## supplemental material for "Post-retrieval noradrenergic activation impairs subsequent memory depending on cortico-hippocampal reactivation"

**Title**

### Supplemental Results

#### Control variables

We controlled for potential group differences in depressive mood, chronic stress as well as state and trait anxiety. Importantly, the three groups did not differ in any of these variables (depressive mood: $F(1,60) = 0.67$, $P = .406$, $\eta^2 = 0.01$, state anxiety: $F(1,60) = 0.31$, $P = .581$, $\eta^2 < 0.01$; trait anxiety: $F(1,60) = 1.07$, $P = .305$, $\eta^2 = 0.02$, chronic stress: $F(1,60) = 0.35$, $P = .557$, $\eta^2 < 0.01$). See Supplemental Table S6.

#### Inter-individual differences in pharmacological responses did not significantly mediate subsequent memory

To account for individual differences in cortisol responses after pill intake, we fit additional GLMMs predicting Day 3 subsequent memory of cued and correct trials including the factors *Individual baseline-to-peak cortisol* and *Group*. Doing so allowed us to account for variation in Day 3 performance, which might have resulted from within-group variation in cortisol responses, in particular in the CORT group. Importantly, none of the models predicting Day 3 memory performance by Day 2 cortisol-increase and Group, median-split RTs (high/low), hippocampal activity and RTs, or hippocampal activity and VTC category reinstatement revealed a significant group x baseline-to-peak cortisol interaction (all Ps > .122). These results suggest that inter-individual differences in cortisol responses did not have a significant impact on subsequent memory, beyond the influence of group per se. The same analyses were repeated for systolic blood pressure employing GLMMs predicting Day 3 subsequent memory of cued and correct trials including the factors *Individual baseline-to-peak systolic blood pressure* and *Group* to account for variation in Day 3 performance, which might have resulted from within-group variation in blood pressure response, in particular in the YOH group. While the model predicting Day 3 memory performance revealed a significant *Individual baseline-to-peak systolic blood pressure* × *Group* × *median-split RTs (high/low)* interaction ($\beta = -0.05 \pm 0.02$, $z = -2.04$, $P = .041$, $R^2_{conditional} = 0.01$), post-hoc slope tests, however, did not show any significant difference between groups (all $P_{Corr} > .329$). The remaining models including hippocampal activity and RTs, or hippocampal activity and VTC category reinstatement did not reveal a significant *Group* × *Individual baseline-to-peak systolic blood pressure* interaction (all Ps > .101). These results suggest that inter-individual differences in systolic blood pressure responses did not have a significant impact on subsequent memory, beyond the influence of group per se.

#### Offline Encoding and Cueing Reinstatement in Hippocampus and Cortical Areas

Aside from examining neural activity related to retrieval during the Memory Cueing task, we also investigated evidence of offline reactivation, which is manifested in neural reinstatement observed during the resting-state scan conducted both before and after the Memory Cueing task. An ANOVA including the factors *Group*, *Time* (pre, post), and *Threshold* (pre, post) revealed a significant *Time* × *Threshold* interaction in the hippocampus ($F(1,58) = 40.48$, $P < .001$, $\eta^2 = 0.02$). Post-hoc t-tests confirmed a significant increase from pre- to post-cueing hippocampal offline reactivation in the pre-threshold condition (post-pre: $t(60) = -3.21$, $P =$

.006, d = 0.59) compared to post-threshold (post-pre: t(60) = -0.78, P = 1, d = 0.05), without a difference between groups (all Ps > .162). In addition to this analysis of the offline reactivation of the memory cueing events, we analyzed the hippocampal offline reactivation of encoding patterns. We again utilized an ANOVA including the factors *Group*, *Time* (pre, post), and *Threshold* (pre, post) which did not reveal any significant main effect or interaction of the factors (all Ps > .823) indicating that patterns from Day 1 memory encoding were not reactivated offline in the hippocampus after Day 2, either before memory cueing or after reactivation during memory cueing. We did however not observe a difference in post-cueing offline reactivation of encoding vs. memory cueing patterns (P = .393). Thus, while we obtained evidence for hippocampal offline reactivation of the Day 2 cueing events, there was no significant evidence for hippocampal offline reactivation of the encoded representation after Day 2 Memory Cueing.

Next, we tested for offline reactivation of patterns from memory cueing in the post- vs. pre-cueing resting state intervals in VTC. An ANOVA including the factors *Group*, *Time* (pre, post), and *Threshold* (pre, post) revealed a significant *Time* × *Threshold* interaction (F(1,58) = 49.42, P < .001, $\eta^2$ = 0.02). Post-hoc t-test confirmed again a significant increase from pre- to post-cueing offline reactivation in the pre-threshold condition (post-pre: t(60) = -2.84, P = .018, d = 0.51) compared to post-threshold condition (post-pre: t(60) = 0.30, P = 1, d = 0.05). Testing for offline reinstatement of Day 1 encoding patterns revealed a similar pattern of results. Again, a significant *Time* × *Threshold* interaction was observed (F(1,58) = 32.63, P < .001, $\eta^2$ = 0.02), indicating generally stronger reactivation of encoding patterns after the memory cueing task. Notably, however, we also observed significant interactions including the factor *Group* (*Group* × *Time*: F(2,58) = 5.58, P = .009, $\eta^2$ = 0.08 ; *Group* × *Threshold*: F(2,58) = 6.61, P = .018, $\eta^2$ = 0.09). Post-hoc t-tests revealed that in the post-cueing interval the PLAC group showed significantly less reactivation events in the VTC compared to the YOH and CORT groups (PLAC vs. YOH: t(28.35) = -2.82, P = .017, d = 0.86; PLAC vs. CORT: t(35.92) = -3.09, P = .007, d = 0.96). We did not observe a difference in post-cueing offline reactivation of encoding vs. memory cueing patterns (P = .393). For the VTC, we obtained evidence suggesting offline reactivation of the Day 2 cueing events as well as the encoded representations following Day 2 Memory Cueing.

Finally, we tested for offline reactivation of patterns from memory cueing in the post- vs. pre-cueing resting state intervals in PCC. An ANOVA including the factors *Group*, *Time* (pre, post) and *Threshold* (pre, post) revealed a significant *Time* × *Threshold* interaction (F(1,58) = 22.94, P < .001, $\eta^2$ = 0.01). A Post-hoc t-test revealed a trend for a difference between pre- and post-cueing offline reactivation in the pre-threshold condition (post-pre: t(60) = -2.80, P = .057, d = 0.43), and no difference in the post-threshold condition (post-pre: t(60) = -0.13, P = 1, d = 0.02). Offline reactivation analysis of encoding patterns revealed a similar pattern of results. Again, a significant *Time* × *Threshold* interaction was observed (F(1,58) = 63.13, P < .001, $\eta^2$ = 0.01), indicating generally more reactivation of Day 1 encoding patterns after the memory cueing task. Post-hoc t-tests revealed a trend-level difference between pre- and post-cueing offline reactivation in the pre-threshold condition (post-pre: t(60) = -2.41, P = .056, d = 0.44), and no significant difference in the post-threshold condition (t(60) = 0.25, P = 1, d = 0.05). While we again found evidence indicating offline reactivation of the Day 2 cueing events in PCC, there was only trend-level evidence supporting PCC offline reactivation of the encoding representations following Day 2 Memory Cueing.

**Day 3 Offline Pattern Reactivation does not Modulate Effects of Post-retrieval Noradrenergic Activation on Subsequent Memory Performance**

Next, we asked whether offline reactivation of the retrieval patterns during Day 2 memory cueing interacted with the effects of post-retrieval YOH and CORT in light of subsequent memory effects. We classified cued and correct (Day 2) trials as strongly or weakly reactivated based on a median-split on region-specific (hippocampus, VTC, PCC) offline pattern reinstatement, allowing us to include the uncued trials in further analyses. We employed a GLMM with *Offline reactivation* (post-pre) and *Group* as predictors of Day 3 associative category hits. This analysis did not yield any main effect or interaction including the factor Group (all Ps > .215) in our ROIs (hippocampus, VTC, PCC), suggesting that offline processing of the memory cueing (retrieval-related) patterns was not affected by the pharmacological manipulation. In the next step, we included the offline reinstatement from the encoding task (Day 1) in the model, predicting Day 3 associative category hits. This GLMM did not reveal any main effect or interaction including the factor Group either (all Ps > .993). Together, Day 2 offline reinstatement of encoding- or retrieval-related patterns did not modulate the impact of the post-retrieval stress system manipulations on subsequent memory.

### Supplemental Methods

#### S1: Stimulus Material

The words of each associative word-picture pair had either negative (mean valence = 3.45, mean arousal = 5.72, mean concreteness = 4.62) or neutral valence (mean valence = 5.06, mean arousal = 2.15, mean concreteness = 4.41). These words were selected from the Leipzig Affective Norms for German[84]. Since there was no significant influence of word valence at the behavioural and neural levels, which may be due to the fact that the arousal evoked by emotional words is typically significantly lower than for pictures or movies, we did not include the factor valence in the analyses reported here.

#### S2: Catch trials

Out of the 164 word-picture pairs presented during encoding, 20 pairs were designated as catch trials for the subsequent cued recall tasks. The selection of word-picture catch trial pairs was counterbalanced in terms of valence (negative/neutral) and category (scene/object). Catch trials served to maintain participants' attention during the cued recall tests and to motivate participants to retrieve the associated picture while seeing the associated word. To further motivate participants to reactivate the associated picture in as much detail as possible when seeing the word cue, participants were informed that correctly answered catch trials would increase their financial compensation. The cued recall tests on Days 1 and 3 included eight catch trials each, while the shorter Day 2 Memory Cueing task included four catch trials. The temporal position of catch trials was distributed within a task, ensuring equal spacing between them. A catch trial was triggered when participants correctly designated the presented word as 'old', 'old/scene' or "old/object'. Upon this choice, either the corresponding or a semantically similar picture probe was displayed on the screen for 0.5 s and participants had to judge whether the probe was the studied associate of the word, responding 'yes' or 'no' within 1 s. Catch trial performance did not differ between groups on any experimental day (all Ps > .603). All catch trials were subsequently excluded from the analyses to prevent potential biases in memory effects due to the re-presentation of correct or semantically similar picture probes together with old words.

#### S3: Tracking offline reactivation

In addition to retrieval-related neural activity during the Memory Cueing task, we also tested for evidence of offline reactivation, reflected in neural reinstatement during the resting-state scan before and after the Memory Cueing task. To do so, we employed an RSA approach previously utilized to analyse offline replay[102], correlating trial/event-specific neural patterns from the Memory Cueing task (Day 2) with patterns from the pre- and post-memory cueing resting-state fMRI scans, separately for the hippocampus, VTC, and PCC. To compare the number of pre- and post-reactivation events, we initially calculated the mean +1.5 SDs across all correlations within the pre-cueing phase *(pre-threshold).* This threshold was then applied to pre- and post-cueing correlation matrices. To validate potential effects, we repeated this approach but applied the threshold (mean +1.5 SDs) from the post-cueing interval *(post-threshold)* to the pre- and post-cueing correlation matrices. Offline reactivation events were then quantified as the absolute number of surviving correlations.

**Supplemental Table S1.** Memory performance and reaction times across experimental days.

| | **PLAC** | **YOH** | **CORT** |
|---|---|---|---|
| Day 1 (cued and correct on Day 2) | | | |
| Hits | 49.72 (1.64) | 43.19 (2.15) | 49.59 (2.82) |
| Dprime | 1.21 (0.07) | 1.01 (0.07) | 1.25 (0.11) |
| Reaction time | 2.29 (0.07) | 2.40 (0.07) | 2.39 (0.07) |
| Day 1 (not cued on Day 2) | | | |
| Hits | 49.38 (1.96) | 41.77 (2.11) | 50.64 (2.64) |
| Dprime | 1.19 (0.09) | 0.96 (0.07) | 1.17 (0.10) |
| Reaction time | 2.27 (0.07) | 2.41 (0.09) | 2.44 (0.07) |
| Day 2 (cued and correct on Day 2) | | | |
| Hits | 68.85 (1.52) | 63.95 (8.56) | 69.70 (13.34) |
| Reaction time | 2.27 (0.63) | 2.31 (0.65) | 2.24 (0.63) |
| Reaction time (fast) | 1.86 (0.05) | 1.92 (0.05) | 1.82 (0.04) |
| Reaction time (slow) | 2.80 (0.06) | 2.89 (0.07) | 2.80 (0.06) |
| Day 3 (cued and correct on Day 2) | | | |
| Hits | 74.32 (1.68) | 65.24 (2.12) | 69.80 (2.52) |
| Dprime | 2.25 (0.11) | 1.78 (0.10) | 2.14 (0.13) |
| Reaction time | 1.98 (0.08) | 2.04 (0.07) | 1.98 (0.07) |
| Day 3 (not cued on Day 2) | | | |
| Hits | 46.71 (1.78) | 36.25 (1.80) | 46.16 (2.39) |
| Dprime | 1.16 (0.09) | 0.74 (0.07) | 1.09 (0.09) |
| Reaction time | 2.29 (0.07) | 2.30 (0.08) | 2.31 (0.07) |

Reaction times relate hits specifically. Data represent means ± SE.

**Supplemental Table S2**. Physiological parameters and mood at baseline across Day 1 and Day 3.

| | | PLAC | YOH | CORT |
|---|---|---|---|---|
| Day 1 | Heart rate (bpm) | 78.85 (2.67) | 75.97 (2.77) | 76.35 (2.52) |
| | Systolic blood pressure (mmHg) | 107.80 (2.80) | 116.47 (2.63) | 114.66 (3.66) |
| | Diastolic blood pressure (mmHg) | 68.32 (1.87) | 71.78 (2.18) | 71.00 (2.29) |
| | cortisol (nmol) | 9.49 (2.02) | 7.16 (1.16) | 10.76 (1.56) |
| | Mood (good/bad) | 34.80 (0.90) | 33.47 (0.98) | 33.70 (0.87) |
| | Tiredness (energized/tired) | 31.05 (1.06) | 30.76 (1.14) | 32.30 (1.01) |
| | Calmity (calm/restless) | 30.50 (1.15) | 30.95 (1.47) | 28.90 (1.07) |
| Day 3 | Heart rate (bpm) | 78.65 (2.46) | 78.28 (2.31) | 78.26 (2.38) |
| | Systolic blood pressure (mmHg) | 112.22 (2.14) | 112.42 (3.26) | 114.04 (3.86) |
| | Diastolic blood pressure (mmHg) | 72.15 (1.48) | 72.66 (1.54) | 68.83 (1.95) |
| | cortisol (nmol) | 8.41 (1.59) | 5.88 (1.12) | 7.32 (1.01) |
| | Mood (good/bad) | 34.30 (1.05) | 34.71 (0.81) | 34.10 (0.91) |
| | Tiredness (energized/tired) | 33.40 (1.05) | 34.09 (0.98) | 32.80 (0.85) |
| | Calmity (calm/restless) | 30.50 (1.15) | 31.00 (1.20) | 30.90 (1.00) |

Subjective and physiological parameters of participants taken at the beginning of Day 1 and Day 2. There were no significant difference in either subjective or physiological stress parameters between groups. Data represent means (±SE).

**Supplemental Table S3.** Significant clusters in the whole-brain analyses of Day 2 memory cueing (correct – incorrect).

| Region | Central coordinates (x,y,z; MNI) | Cluster-P (FWE 0.05) | Cluster-T |
|---|---|---|---|
| Frontal Medial, ACC | -4, 50 , -8 | < .001 | 10.72 |
| Lateral Occipital L, AG L | -48, -66, 30 | < .001 | 9.08 |
| Amygdala L, Striatum L | -14, 4, -16 | < .001 | 8.22 |
| Posterior Cingulate Cortex | 4, -41, 38 | < .001 | 8.10 |
| Supramarginal Gyrus R | 62, -44, 20 | < .001 | 7.94 |
| hippocampus L | -26, -32, -10 | < .001 | 7.93 |
| hippocampus R | 32, -40, -12 | < .001 | 7.89 |
| Frontal Pole | -22, 38, 42 | < .001 | 7.32 |
| Temporooccipital Cortex R | 52, -50, -14 | < .001 | 6.13 |
| Superior Parietal R | 30, -46, 62 | < .001 | 6.11 |
| Temporooccipital Cortex L | -52, -62, -2 | < .001 | 6.93 |
| Postcentral Gyrus | 50, -22, 50 | < .001 | 5.77 |
| Superior Frontal Gyrus | -20, 2, 56 | < .001 | 6.67 |
| Putamen L | -26, -8, 8 | < .001 | 6.64 |
| Mid temporal gyrus L | -62, -16, -8 | < .001 | 6.37 |

Depicted clusters are ordered by cluster-peak T-values.

**Supplemental Table S4**. Results from Day 2 gPPI analysis (remembered – forgotten).

| Seed | Target | Cluster size | pFWE (cluster) | pFWE (peak) |
|---|---|---|---|---|
| PCC | Lat. OCC left | 237 | < .001 | .015 |
| | VTC | 1218 | < .001 | < .001 |
| MPFC | Lat. OCC left | 404 | < .001 | < .001 |
| | Sup. Parietal lobe | 836 | < .001 | < .001 |
| | VTC | 1168 | < .001 | .002 |
| Left Hippocampus | VTC | 260 | < .001 | .016 |
| VTC | Lat. OCC left | 789 | < .001 | < .001 |
| | Sup. Parietal lobe | 1123 | < .001 | < .001 |
| | VTC | 260 | < .001 | .016 |
| PCC | Lat. OCC left | 237 | < .001 | .015 |
| | VTC | 1218 | < .001 | < .001 |
| MPFC | Lat. OCC left | 404 | < .001 | < .001 |
| | Sup. Parietal lobe | 836 | < .001 | < .001 |
| | VTC | 1168 | < .001 | .002 |
| Left Hippocampus | VTC | 260 | < .001 | .016 |
| VTC | Lat. OCC left | 789 | < .001 | < .001 |
| | Sup. Parietal lobe | 1123 | < .001 | < .001 |

All P-values are Bonferroni-corrected for amount of ROIs. Only clusters with a minimum of 25 voxels were included.

**Supplemental Table S5**. Physiological parameters across Day 2.

| | | **PLAC** | **YOH** | **CORT** |
|---|---|---|---|---|
| Heart rate (bpm) | Base | 80.42 (2.41) | 75.02 (2.10) | 77.52 (1.97) |
| | Task | 87.40 (2.29) | 81.61 (2.06) | 82.76 (1.86) |
| | Resting state post | 80.15 (2.28) | 74.42 (2.21) | 75.28 (2.18) |
| | +40 | 70.35 (2.58) | 64.92 (2.06) | 67.04 (2.10) |
| | +55 | 70.65 (2.28) | 65.59 (1.97) | 67.52 (2.33) |
| | +70 | 69.12 (2.18) | 67.45 (2.37) | 65.61 (2.03) |
| | +85 | 68.90 (2.40) | 67.47 (1.97) | 65.73 (2.12) |
| | +100 | 70.25 (2.78) | 69.45 (2.60) | 67.85 (2.16) |
| Systolic BP (mmHg) | Base | 113.00 (3.11) | 115.42 (3.11) | 117.95 (3.23) |
| | +40 | 115.35 (1.95) | 118.00 (3.27) | 117.38 (3.70) |
| | +55 | 110.40 (1.75) | 118.61 (3.36) | 115.83 (3.11) |
| | +70 | 108.45 (1.71) | 120.92 (3.43) | 115.11 (3.68) |
| | +85 | 108.50 (2.27) | 122.38 (3.57) | 115.28 (3.34) |
| | +100 | 107.55 (2.46) | 123.69 (3.29) | 115.78 (3.21) |
| Diastolic BP (mmHg) | Base | 72.53 (1.99) | 71.86 (2.61) | 74.57 (2.14) |
| | +40 | 73.22 (2.26) | 74.14 (1.96) | 73.76 (2.08) |
| | +55 | 72.33 (1.45) | 71.02 (2.10) | 72.67 (2.05) |
| | +70 | 71.28 (1.64) | 73.31 (2.36) | 72.98 (2.18) |
| | +85 | 72.85 (1.67) | 75.79 (2.12) | 73.31 (2.30) |
| | +100 | 71.55 (1.71) | 77.29 (2.30) | 73.98 (1.81) |
| cortisol (nmol) | Base | 11.14 (2.14) | 7.22 (1.17) | 8.18 (1.13) |
| | +20 | 5.40 (0.78) | 4.04 (0.39) | 8.15 (2.03) |
| | +40 | 4.53 (0.62) | 2.92 (0.31) | 24.59 (6.32) |
| | +70 | 3.63 (0.60) | 3.14 (0.41) | 57.60 (4.88) |
| | 100 | 2.70 (0.34) | 4.05 (0.69) | 52.93 (4.95) |
| Skin conductance | Resting state pre | 5.82 (1.35) | 3.32 (0.55) | 5.97 (0.86) |
| | Task | 5.88 (1.47) | 3.07 (0.75) | 6.71 (1.31) |
| | Resting state post | 9.28 (1.63) | 6.53 (0.96) | 10.02 (1.25) |

Physiological parameters of participants across Day 2 relative to drug administration. Systolic blood pressure significantly increased over time in the YOH group, compared to the CORT and PLAC group. Salivary cortisol increased in the CORT group but not in the YOH or PLAC group. Data represent means (±SE).

**Supplemental Table S6.** Subjective mood scores across Day 2.

| | | PLAC | YOH | CORT |
|---|---|---|---|---|
| MDBF *Base* | Mood (good/bad) | 34.85 (1.11) | 34.38 (0.78) | 34.90 (0.72) |
| | Tiredness (energized/tired) | 33.85 (1.10) | 33.33 (1.04) | 33.70 (1.03) |
| | Calmness (calm/restless) | 34.85 (1.11) | 34.38 (0.78) | 34.90 (0.72) |
| MDBF *+40* | Mood (good/bad) | 35.05 (1.08) | 34.81 (0.81) | 33.70 (0.81) |
| | Tiredness (energized/tired) | 33.70 (0.94) | 33.53 (1.06) | 32.95 (1.01) |
| | Calmness (calm/restless) | 30.55 (0.87) | 28.90 (1.47) | 30.04 (0.91) |
| MDBF *+55* | Mood (good/bad) | 34.70 (0.86) | 35.14 (0.80) | 33.20 (1.09) |
| | Tiredness (energized/tired) | 34.10 (0.98) | 33.80 (0.92) | 32.90 (1.10) |
| | Calmness (calm/restless) | 30.00 (0.97) | 30.47 (1.30) | 28.45 (1.75) |
| MDBF *+70* | Mood (good/bad) | 34.55 (1.11) | 34.81 (0.86) | 32.75 (1.07) |
| | Tiredness (energized/tired) | 33.65 (1.08) | 34.28 (1.08) | 32.75 (1.16) |
| | Calmness (calm/restless) | 29.55 (1.43) | 30.81 (1.34) | 29.60 (1.38) |
| MDBF *+85* | Mood (good/bad) | 34.20 (1.14) | 34.09 (0.93) | 32.30 (1.20) |
| | Tiredness (energized/tired) | 33.20 (1.18) | 33.47 (1.23) | 31.60 (1.14) |
| | Calmness (calm/restless) | 28.80 (1.73) | 30.71 (1.56) | 28.55 (1.58) |
| MDBF *+100* | Mood (good/bad) | 33.30 (1.39) | 33.19 (1.16) | 32.55 (1.02) |
| | Tiredness (energized/tired) | 32.70 (1.34) | 32.61 (1.38) | 32.50 (1.26) |
| | Calmness (calm/restless) | 30.20 (1.36) | 29.19 (1.56) | 29.40 (1.48) |

Subjective mood ratings according to the *Mehrdimensionale Befindlichkeitsfragebogen* (MDBF[85]) across day 2. Scores did not reveal a significant difference in either scale between groups. Data represent means (±SE).

**Supplemental Table S7**. Participants' state, trait anxiety, chronic stress and depression scores.

| | PLAC | YOH | CORT |
|---|---|---|---|
| Depression score | 6.20 (0.89) | 9.04 (1.20) | 7.47 (1.00) |
| State anxiety | 43.20 (1.02) | 42.09 (0.94) | 42.38 (1.16) |
| Trait anxiety | 43.05 (1.41) | 46.95 (1.79) | 45.19 (0.93) |
| Chronic stress | 70.95 (6.86) | 90.61 (8.42) | 77.14 (5.37) |

State and Trait anxiety scores were measured with the State-Trait Anxiety Inventory. Depression Scores were determined utilizing the Beck Depression Inventory. Chronic stress was measured with the Trier Inventory of Chronic Stress. Participants conducted the three questionnaires at home before the actual experiment started. Data represent means (±SE).
